# Two Isoforms of the Guanine Nucleotide Exchange Factor, Daple/CCDC88C Cooperate as Tumor Suppressors

**DOI:** 10.1101/312934

**Authors:** Ying Dunkel, Jason Ear, Yash Mittal, Blaze B. C. Lim, Lawrence Liu, Magda K. Holda, Ulrich Nitsche, Jorge Barbazán, Ajay Goel, Klaus-Peter Janssen, Nicolas Aznar, Pradipta Ghosh

## Abstract

Previously Aznar et al., showed that Daple enables Wnt/Frizzled receptors to transactivate trimeric G proteins during non-canonical Wnt signaling via a novel G-protein binding and activating (GBA) motif. By doing so, Daple serves as a double-edged sword; earlier during oncogenesis it suppresses neoplastic transformation and tumor growth, but later it triggers epithelial messenchymal transition (EMT). We have identified and characterized two isoforms of the human Daple/CCDC88c gene. While both isoforms cooperatively suppress tumor growth via their GBA motif, only the full-length transcript triggers EMT and invasion. Aspirin suppresses the full-length transcript and protein but upregulates the short isoform. Both isoforms are suppressed during colon cancer progression, and their reduced expression carries additive prognostic significance. These findings provide insights into the opposing roles of Daple during cancer progression and define the G protein regulatory GBA motif as one of the minimal modules essential for Daple’s role as a tumor suppressor.

## Introduction

Earlier we defined a novel paradigm in Wnt signaling in which Frizzled receptors (FZDRs) activate the G proteins and trigger non-canonical Wnt signaling via Daple (CCDC88C) (<u>Aznar et al., 2015a</u>). Daple, a multimodular signal transducer and a cytosolic protein was originally discovered as a Dishevelled (Dvl)-binding protein that regulates Wnt signaling (<u>Kobayashi et al., 2005</u>; <u>Oshita et al., 2003</u>). Subsequent work showed that Daple directly binds ligand-activated FZDs, and serves as a guanine-nucleotide exchange factor (GEF) that activates the G protein, Gαi (<u>Aznar et al., 2015a</u>). Binding to the G protein is mediated via Daple’s C-terminally located Gα-binding and activating (GBA) motif; such binding triggers non-canonical activation of Gαi (<u>Aznar et al., 2015a</u>). Binding to FZDR is also brought on via a C-terminally located stretch distal to the GBA motif. Upon ligand stimulation, Daple-GEF dissociates from Dvl, binds and displaces Dvl from FZDs, and assembles Daple-Gαi complexes. How Daple:Dvl complexes are disassembled was unknown until recently when we demonstrated that phosphorylation of Daple’s PDZ-binding motif (PBM) by both receptor and non-receptor tyrosine kinases can trigger this change (<u>Aznar et al., 2018</u>). Disassembly of Daple:Dvl complexes and formation of FZD:Daple:Gαi complexes facilitates the activation of trimeric Gαi near ligand-activated FZDs. Daple activates Gαi within the FZD:Daple:Gαi complexes; such non-canonical activation of Gαi by Daple-GEF suppresses cAMP, whereas released ‘free’ Gβγ heterodimers enhance Rac1 and PI3K-Akt signals (<u>Aznar et al., 2015a</u>). Although Daple-dependent enhancement of non-canonical Wnt signals can suppress tumor growth (<u>Aznar et al., 2015a</u>), it can also fuel EMT, trigger cancer cell migration and invasion (<u>Aznar et al., 2015a</u>; <u>Ishida-Takagishi et al., 2012</u>) and drive metastasis (<u>Niavarani et al., 2016</u>). Furthermore, elevated expression of Daple-GEF in circulating tumor cells prognosticates a poor outcome (<u>Barbazan et al., 2016</u>). In doing so, Daple behaves like a double-edged sword--a tumor suppressor early during oncogenesis which mimics an oncogene and fuels metastatic invasion later.

Here we identified a novel Daple isoform (Daple-V2) which corresponds only to the C-terminal region of full length Daple (Daple-fl; RefSeq). We demonstrate that Daple-V2 possesses all the functional domains to bind Dvl, FZDR and Gαi and represents the smallest autonomous Daple unit able to inhibit the β-catenin/TCF/LEF pathway and suppress tumor cell growth. We also demonstrate that, both isoforms have different subcellular distribution, and unlike Daple-fl, Daple-V2 does not enhance tumor cell invasiveness and therefore, is a more potent tumor suppressor and a better prognostic marker.

## Results and Discussion

### Daple-V2 represents the smallest autonomous Daple unit able to bind Dvl, FZD7R and Gαi

We noted that there are 5 other transcript variants catalogued in Ensembl and UniProt databases, all predicted to code proteins: V2 (552aa), V3 (502aa), V4 (478aa), V5 (96aa) and V6 (88aa) (**Fig 1A**). Among them, V2, V3 and V4 are predicted to encode stretches within Daple’s C-terminus, whereas V5 and V6 are predicted to encode stretches within Daple’s N-terminus. Because distinct protein isoforms generated from single genes are known to contribute to the diversity of the proteome (<u>Larochelle, 2016</u>), we asked if Daple’s seemingly opposing and bifaceted roles in cancers may be, in part, due to the functions of two isoforms of the same protein. We focused on the 2 variants of Daple transcript (Daple-V2 and -V3) (**Fig 1**); both represent the last 5 exons of Daple-fl and, if translated, we noted that both isoforms should contain both a functional GBA motif and a C-terminal PBM motif **(Fig 1A, B; Fig 1-Figure Supplement 1A-B)**. We noted that Daple-V2 and V3 differs from Daple-fl by a unique 5’ end **(Fig 1B; Fig 1-Figure Supplement 1B)** which adds a unique sequence comprised of 5 amino acids (MSVLS) on the N-terminus of the isoform. To analyze if either of the transcripts are expressed, first we amplified Daple cDNA from HeLa cells using reverse transcription-PCR (RT-PCR) with specific Daple primers that can detect Daple-V2 (552aa) and -V3 (506aa) short isoforms (see *Materials and Methods*). Indeed two new transcripts were amplified (see **Fig 1-Figure Supplement 2**). By cloning the products into pcDNA3.1 vector and sequence analyses we confirmed that the most abundant isoform in human cells and colonic tissues is Daple-V2.

**Figure 1:**
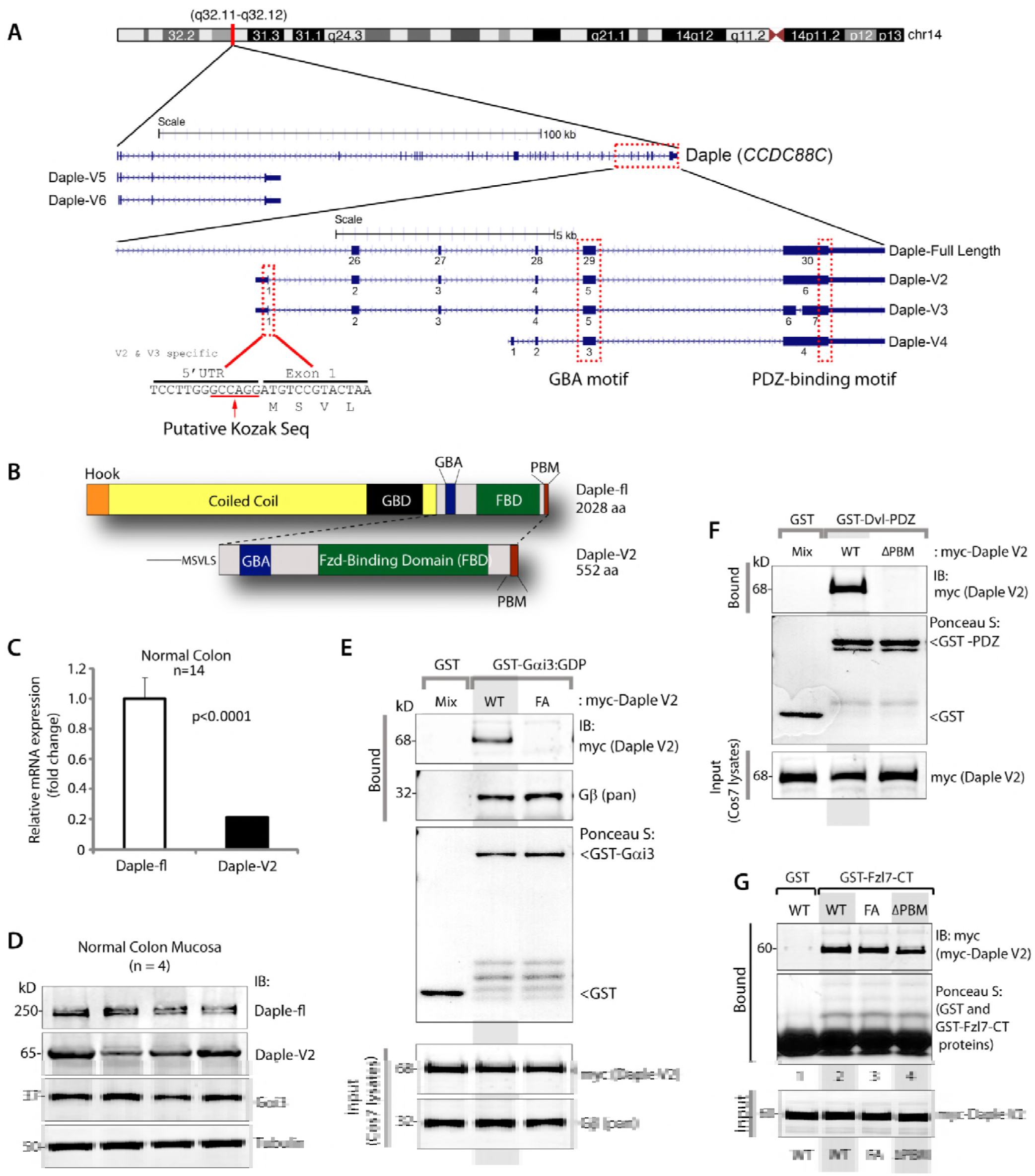
Identification and characterization of a short isoform of CCDC88C (Daple-V2) that contains minimal C-terminal modules to bind trimeric Gαi, Dvl and Frizzled receptor. **(A)** Schematic showing the various N-terminal and C-terminal transcripts of Daple. Various isoforms containing the unique modular C-terminus of the protein are highlighted. All C-terminal transcripts contain an exon coding for a GBA motif and a PDZ-Binding motif. Daple-V2 and Daple-V3 transcripts contain an unique 5’ end that is transcribed from an intronic region between exon 25 and 26 of the full length gene. This 5’ end contains exon 1 of Daple-V2 and Daple-V3 and codes for an unique N-terminal peptide on these isofrms. The 5’ UTR of the two isoform contains a putative kozak sequence. Names of the RNA transcript and encoded protein is indicated on the right. **(B)** Schematic comparing the domain distribution of Daple-fl and the shorter isoform Daple-V2. **(C)** mRNA isolated from 14 normal colon samples were analyzed for the expression of full length (fl) or short (V2) isoform of Daple. Relative mRNA expression (Y axis) of both isoforms is displayed as bar graphs. **(D)** Whole cell lysates of colonic epithelial from normal subjects were analyzed for Daple, Gαi3 and tubulin by immunoblotting (IB). Both full length (fl) and short isoform (V2) were detected. **(E)** Purified, recombinant GST-Gαi3 preloaded with GDP and immobilized on glutathione-agarose beads was incubated with cell lysates of Cos7 cells (input) expressing myc-Daple-V2 WT or F194A (FA) as indicated. Bound proteins were analyzed for Daple-V2 (myc) and Gβ by immunoblotting (IB). Equal loading of GST-tagged proteins were confirmed by Ponceau S staining. F194A mutation disrupts binding of Daple-V2 to Gαi3. **(F)** Purified, recombinant GST-tagged PDZ domain of Dvl immobilized on glutathione-sepharose beads was incubated with cell lysates of Cos7 cells (input) expressing myc-Daple-V2 WT or delta PBM (ΔPBM) as indicated. Bound proteins were analyzed for Daple-V2 (myc) by immunoblotting (IB). Equal loading of GST-tagged proteins was confirmed by Ponceau S staining. Deletion of the C-terminal PDZ-binding motif disrupts binding of Daple-V2 to PDZ domain of Dvl. **(G)** Purified, recombinant GST-tagged carboxy terminus of FZD7R (Fzl7-CT) immobilized on glutathione-sepharose beads was incubated with cell lysates of Cos7 cells (input) expressing myc-Daple-V2 WT, FA or delta PBM (ΔPBM) as indicated. Bound proteins were analyzed for Daple-V2 (myc) by immunoblotting (IB). Equal loading of GST-tagged proteins was confirmed by Ponceau S staining. WT and mutants of Daple-V2 bound similarly to FZD7R.

Because Daple expression is dysregulated during the progression of colon cancer (<u>Aznar et al., 2015a</u>), we first asked if mRNA and protein for both isoforms are expressed in the normal colon, and if so, what might be their relative abundance in normal colon. When we analyzed the copy numbers of Daple-V2 mRNA in 14 colon samples by qPCR, we found the relative abundance of Daple-V2 to be ~20% of that of Daple-fl (**Fig 1C**). We also confirmed the expression of Daple-V2 protein (**Fig 1D**) by analyzing lysates of mucosal biopsies taken from normal colons by immunoblotting using an anti-Daple-CT antibody raised against aa 1660-2028 [previously validated in (<u>Aznar et al., 2015a</u>)] and expected to detect both Daple-fl and -V2 isoforms].

To study the properties of Daple-V2 and compare them with the previously described full-length protein, we cloned the Daple-V2 transcript into a myc-pcDNA vector plasmid for mammalian expression as a N-terminally myc-tagged protein, just as we did previously for Daple-fl (<u>Aznar et al., 2015a</u>). As expected, we confirmed by GST pulldown assays that myc-Daple-V2 interacts with Gαi3, PDZ domain of Dvl and the cytoplasmic tail of FZDR7 **(Fig 1E-G).** These findings indicate that Daple-V2 represents the smallest autonomous Daple unit possessing all the biochemical features of Daple-fl previously described (<u>Aznar et al., 2015a</u>).

### Daple-V2 antagonizes Wnt signaling via the β-Catenin/TCF/LEF pathway, suppresses growth and proliferation but does not trigger EMT or cell invasion

We previously demonstrated that Daple-fl and more specifically its GBA motif, antagonizes the β-catenin-dependent Wnt signaling pathway and inhibits colony growth, but enhances the PI3K-Akt and Rac1 signals, EMT and invasion (<u>Aznar et al., 2015a</u>). Because Daple-V2 possesses a GBA motif, and because the motif is required for binding G proteins, we asked if this motif is functional in cells. Using DLD1 colon cancer cells stably 7-TGP **(Fig 2A)**, an eGFP expressing Wnt activity reporter construct, or parental DLD1 cells (**Fig 2B**), we generated stable cell lines expressing Daple-V2 wild-type (WT) and GEF-deficient (F194A; FA) mutant. For comparison, we used previously developed and characterized (<u>Aznar et al., 2015a</u>) DLD1 lines expressing the WT Daple-fl.

**Figure 2:**
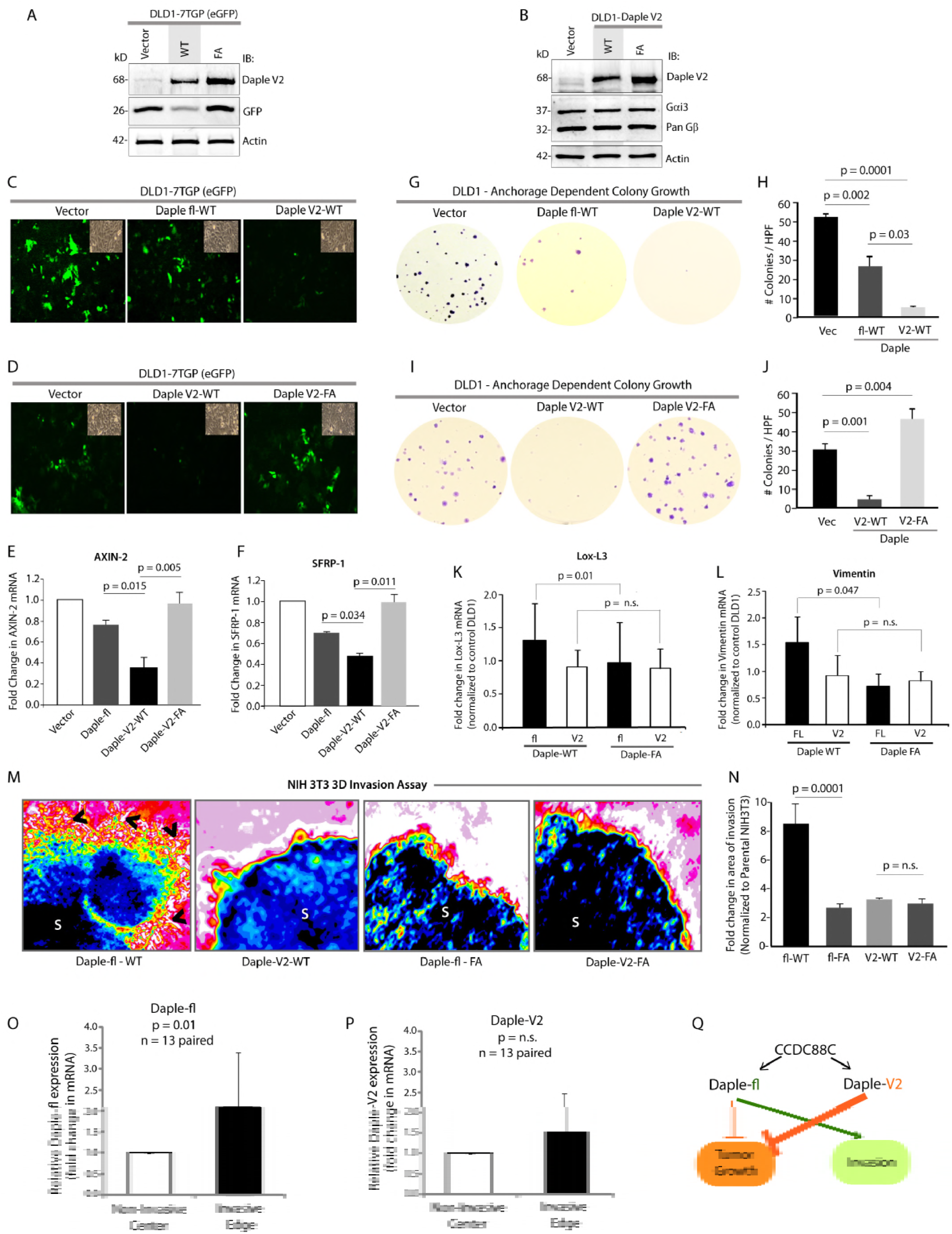
Daple-V2 is a potent suppressor of the β-Catenin/TCF/LEF pathway and tumor growth but has no effect on EMT or cell invasion. **(A)** Whole cell lysates of DLD1 cells stably co-expressing the 7TGP reporter and either vector control or Daple-V2 were analyzed for Daple-V2, GFP and actin by immunoblotting (IB). The intensity of GFP indicates the extent of β-Catenin/TCF/LEF signals. **(B)** Whole cell lysates of DLD1 cells stably expressing vector control, Daple-V2 WT or FA were analyzed for Daple-V2, Gαi3, pan Gβ and actin by immunoblotting (IB). **(C-D)** Monolayers of DLD1 7TGP cell lines in **A** were starved and stimulated with Wnt5a. Images display representative fields analyzed by fluorescence microscopy. The intensity of eGFP signals denotes Wnt transcriptional activity. Insets show representative fields confirming the confluency of the monolayers in each case. In **C**, compared to DLD1 cells expressing vector control, both Daple-fl-WT and Daple-V2-WT showed inhibition of eGFP; inhibition with Daple-V2 was more robust. In **D**, Daple-V2-WT, but not Daple-V2-FA inhibited eGFP. **(E-F)** HeLa cells transfected with myc-Daple constructs as indicated were analyzed for AXIN-2 and SFRP-1 mRNA by qPCR. Results were normalized internally to mRNA levels of the housekeeping gene, GAPDH. Bar graphs display the fold change in each RNA (Y axis) normalized to the expression in cells expressing control vector. Error bars represent mean ± S.D of 3 independent experiments. As shown in the case of Daple-fl previously (Aznar et al., eLife 2015), the GBA motif of Daple-V2 is required for suppression of Wnt target genes. **(G-J)** Monolayers of DLD1 cells in **B** were analyzed for their ability to form adherent tumor cell colonies on plastic plates during 2-3 weeks prior to fixation and staining with crystal violet. In panels **G** and **I** photographs of a representative well of the crystal violet-stained 6-well plates are displayed. The number of colonies was counted by ImageJ (Colony counter). In panels **H** and **J** bar graphs display the # of colonies per well (Y axis) seen in each cell line in **G** and **I**, respectively. Panels **G-H** show that both Daple-fl and Daple-V2 can inhibit tumor growth; the latter is more efficient that the former. Panels **I-J** show that the GBA motif of Daple-V2 is required for the inhibition of anchorage-dependent colony growth. **(K-L)** mRNA expression of the EMT markers LOX-L3 and Vimentin were analyzed by qPCR. Results were normalized internally to mRNA levels of the housekeeping gene, GAPDH. Bar graphs display the fold change in each RNA (Y axis) normalized to the expression in cells expressing vector control. Error bars represent mean ± S.E.M of 3 independent experiments. Daple-fl, but not Daple-V2 enhances the expression of genes that trigger EMT, and such enhancement requires an intact GBA motif. **(M-N)** Spheroids (S) of NIH3T3 cells expressing WT or FA mutant of myc-Daple-fl or myc-Daple-V2 isoform were analyzed for their ability to invade matrigel in response to Wnt5a using a Cultrex-3D Spheroid Invasion Kit (Trevigen). Representative images of spheroid edges are displayed (**M**). An increase of invasion tracks (arrowheads) was noted only from the edge of tumor spheroids formed by cells expressing myc-Daple-fl-WT, but not the GEF-deficient F1675A (FA) mutant. Neither the WT nor the FA mutant of Daple-V2 could trigger invasion. Area of invasion was quantified using ImageJ and displayed as bar graphs (**N**). Error bars representing mean ± S.D of 3 independent experiments. **(O-P)** Paired samples from non-invasive center and the invasive edges of colorectal cancers were analyzed for Daple-fl (**O**) and Daple-V2 (**P**) expression by qPCR. Bar graph displays the relative abundance of Daple expression (Y axis). Daple-fl, but not Daple-V2 is increased in the invading margins of tumors compared to the non-invasive tumor cores. Error bars represent mean ± S.D. n = 13. **(Q)** Schematic summarizing the effect of the newly discovered GBA motif in Daple-fl and Daple-V2 on tumor growth and tumor invasion. The GBA motif of Daple-fl inhibits tumor growth and enhances tumor invasion, whereas the GBA motif of Daple-V2 exclusively inhibits tumor growth. Red lines = Inhibition. Green lines = Enhancement. Thickness of the lines depicts the relative strength of phenotypes.

We found that Daple-V2 and Daple-fl have two similarities and one dissimilarity. First similarity was that both Daple-fl and Daple-V2 antagonize the β-catenin/TCF/LEF pathway **(Fig 2C)**; Daple-V2-WT, but not Daple-V2-FA failed to inhibit eGFP expression (**Fig 2A, D**), indicating that the inhibitory effect of Daple-V2 on the canonical Wnt pathway required an intact GBA motif. Consistently, both Daple-fl and Daple-V2 reduced the transcription of downstream target genes Axin-2 and SFRP-1; once again, the presence of an intact GBA motif was critical for such inhibition (**Fig 2E-F**). Second similarity was that expression of either Daple-fl or Daple-V2 inhibited anchorage-dependent colony growth of DLD1 cells by ~50% and 90% respectively (**Fig 2G-H)**. Such growth suppressive effect required an intact GBA motif because, compared to Daple-V2 WT, expression of the GBA-deficient F1675A (<u>Aznar et al., 2015a</u>; henceforth, FA) mutant not only failed to inhibit cell colony formation, but also enhanced anchorage-dependent growth **(Fig 2I-J)**.

The dissimilarity between Daple-fl and Daple-V2 was observed in their ability to trigger EMT--compared to cells expressing Daple-V2 or Daple-fl FA, those expressing Daple-fl WT had significantly higher expression of Lox-L3 and Vimentin, two genes commonly associated with epithelial-mesenchymal transition (EMT) **(Fig 2K-L).** Furthermore, in 3-D matrigel invasion assays using the transformed NIH3T3 cells exactly as done previously (<u>Aznar et al., 2015a</u>). Enhanced invasion, as determined by the area of invasion was detected exclusively in the presence of Daple-fl WT, but not in cells expressing Daple-V2 or Daple-fl FA, indicating that only Daple-fl can trigger cell invasion (**Fig. 2M-N**). We found that expression of Daple-fl, but not Daple-V2 is increased in the invading margins of tumors compared to the non-invasive tumor cores **(Fig 2O-P)**, which is in keeping with our findings that Daple-V2 does not contribute to EMT or higher invasiveness. These findings demonstrate that compared to Daple-fl, Daple-V2 serves as a more potent inhibitor of the β-Catenin-dependent canonical Wnt pathway and tumor growth in colonies, but it does not enhance EMT or invasion **(Fig 2Q)**.

### The anti-proliferative roles of Daple-fl and Daple-V2 are additive; simultaneous suppression of both transcripts in colon cancers carries poor prognosis

Because both Daple-fl and Daple-V2 suppress colony growth, we asked if such effects are additive. To investigate this, we carried out growth curve assessment and cell viability assays on HeLa cells lines that have been depleted of endogenous Daple by shRNA and stably expressing myc-Daple-fl or myc-Daple-V2 either alone, or together [**Fig 3A**; Daple-depleted HeLa lines have been extensively characterized using a variety of approaches (<u>Aznar et al., 2015a</u>)]. In both assays we found that co-expression of Daple-fl and Daple-V2 isoforms suppressed cell growth and proliferation most effectively compared to either isoforms alone (**Fig 3B-C**).

**Figure 3:**
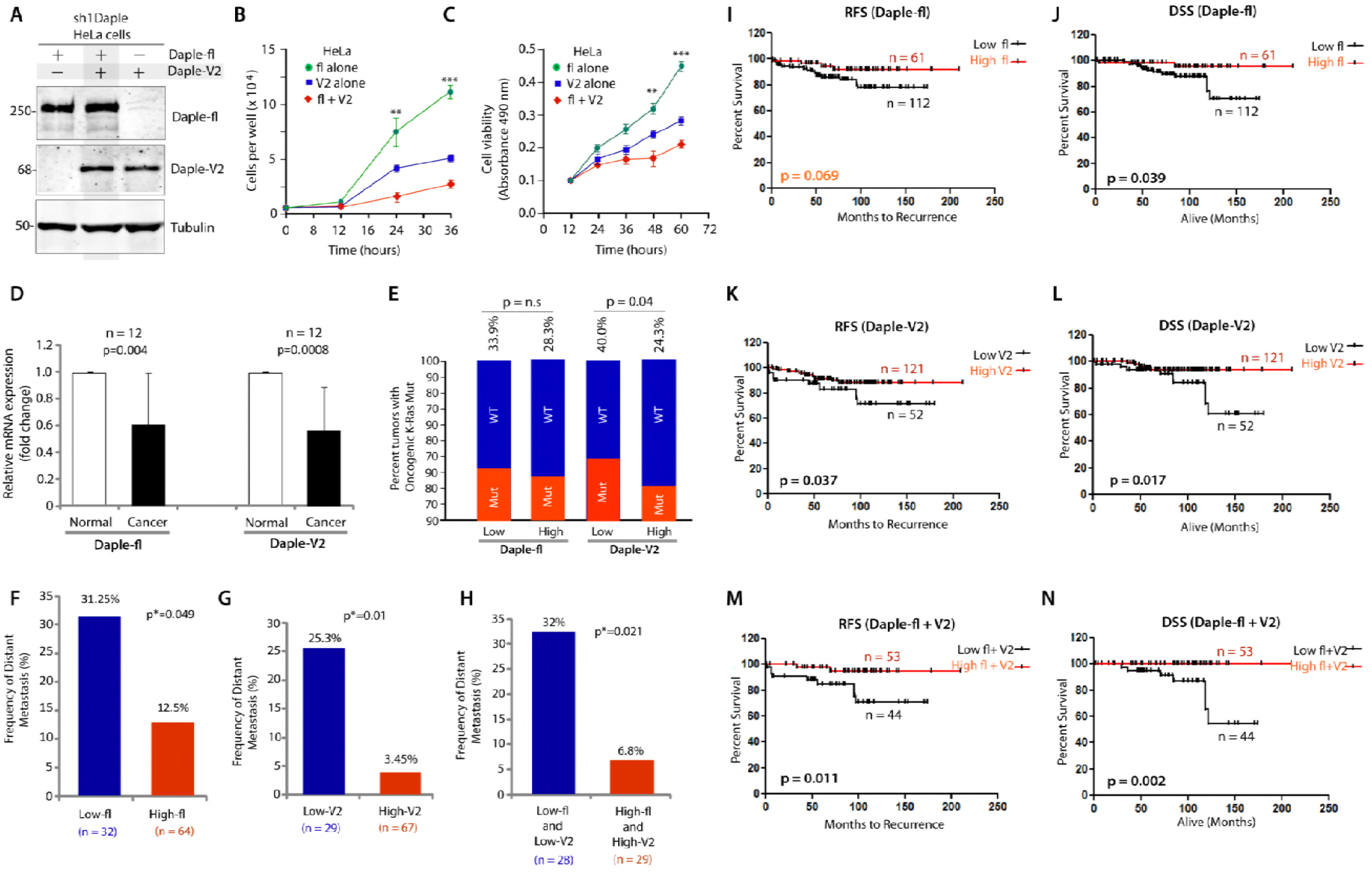
The full length (Daple-fl) and short (Daple-V2) isoforms of Daple cooperatively suppress cell proliferation and their low expression in stage II colorectal cancers carries a worse prognosis. **(A-C)** HeLa cells depleted of Daple (sh1Daple) stably expressing either Daple-fl alone, or Daple-V2 alone or both were analyzed for Daple expression by immunoblotting **(A)** and rate of cell proliferation assays **(B, C)**. Graphs display the rates of proliferation of various HeLa-GIV cell lines, as determined by cell counting (**B**) and cell viability assays (**C**). Results are presented as mean ± S.E.M; n = 3. ** *p* < 0.01; *** p <0.001. **(D)** Paired colorectal tumors and their adjacent normal tissue were analyzed for relative expression of Daple isoforms by qPCR. Bar graph displays the relative abundance of Daple expression (Y axis). Error bars represent mean ± S.D. **(E)** 173 stage II colorectal cancers with known K-Ras mutant status were analyzed for levels of expression of Daple-fl and Daple-V2 mRNA by Taqman qPCR and normalized to GAPDH. Optimal cut-off values for Daple mRNA expression were statistically derived (see detailed *“Materials and Methods”*) to generate subgroups of patients with high or low expression levels. The number of tumors with or without mutant K-Ras that had either low or high expression of Daple isoforms is tabulated in **Figure 3-source data 1**. Bar graphs display the incidence (expressed as %) of K-Ras mutation (Y axis) when either Daple isoforms are either high or low. Red and blue colors indicate whether the tumors harbored oncogenic mutant or WT copy of K-Ras, respectively. The incidence of mutation is displayed on the top of each bar. Tumors with low Daple-V2 had a significant chance that they also harbor mutant K-Ras. No such relationship was seen between levels of expression of Daple-fl and mutant K-Ras. **(F-H)** Bar graphs display the incidence of distant metastasis (as %; Y axis) in stage II colorectal cancers with either low or high levels of expression of Daple-fl alone (**F**), or Daple-V2 alone (**G**), or both Daple isoforms (**H**). **(I-N)** Kaplan-Meier plot of recurrence-free (RFS) and disease-specific (DSS) survival curves of patients with stage II colorectal cancer are stratified by their levels of expression of Daple-fl alone (**I-J**), or Daple-V2 alone (**K-L**), or both Daple isoforms (**M-N**). In the RFS curves, cancers with low Daple-V2 alone exhibited decreased recurrence-free survival (**K**; significant by Log-Rank test). Although a similar trend was seen also in the case of Daple-fl (**I**), significance was not reached. Cancers with low levels of both isoforms exhibited decreased recurrence (**M**) with higher significance than each isoform alone. In the DSS curves, cancers with low Daple-fl alone or Daple-V2 alone exhibited decreased disease specific survival (**J, L**; significant by Log-Rank test). Cancers with low levels of both isoforms exhibited decreased survival (**N**) with higher significance than each isoform alone.

We previously showed that Daple-fl is downregulated earlier during cancer initiation (at the stage of polyp to cancer conversion), and that its expression at high levels carries a good prognosis in colorectal cancers (CRCs) (<u>Aznar et al., 2015a</u>). We also showed that levels of Daple-fl is increased later during metastatic progression and in circulating tumor cells (CTCs), and that its expression at high levels carries a worse prognosis. We asked if and how the expression of Daple-V2 changes during cancer progression in the colon and what, if any, may the prognostic impact of such changes. Once again, we found several similarities and one dissimilarity. Analysis by qPCR in 12 paired colorectal tumors and their adjacent normal tissue showed that, much like Daple-fl, the expression of Daple-V2 is decreased in CRCs **(Fig 3D)**. When we analyzed a cohort of patients with Duke’s Stage II CRCs, we found that tumors that express low Daple-V2 had a significantly higher incidence of oncogenic K-ras mutation **(Fig 3E; Table 1)**; no such correlation was seen in the case of Daple-fl. Tumors that express low Daple-fl (**Fig 3F**) or low Daple-V2 (**Fig 3G**) or low levels of both isoforms **(Fig 3H)** have a higher frequency of progression to distant metastases. Kaplan-Meier analyses revealed that Daple-V2 is a better prognosticator than Daple-fl (compare **Fig 3I-J** to **3K-L**) when used standalone to stratify risk for recurrence-free survival (RFS) and disease-specific survival (DSS). When accounting for high vs low levels of both Daple isoforms, an additive prognostic impact was seen compared to each alone (**Fig 3M-N**). A correlation analysis showed that Daple-V2, but not Daple-fl negatively correlates with the marker for proliferative index of tumor cells Ki67 (<u>Ellis et al., 2017</u>; <u>Niikura et al., 2012</u>), and positive correlation with the tumor suppressor SAM and SH3-Domain Containing 1 ((<u>Martini et al., 2011</u>; <u>Zeller et al., 2003</u>)) (**Table 2**). Together with our findings on cell lines (growth curve, Wnt reporter and tumor cell colony formation assays) these analyses on patient tumors suggest that while both isoforms suppress tumor cell proliferation, Daple-V2 may be a more potent suppressor of tumor cell growth and proliferation than Daple-fl. These findings also define the profile of dysregulated Daple-fl and Daple-V2 expression during oncogenic progression in the colon: both isoforms are suppressed during colorectal cancer progression, and low expression levels of both isoforms alone or simultaneously exhibit decreased survival.

**Table 1.**
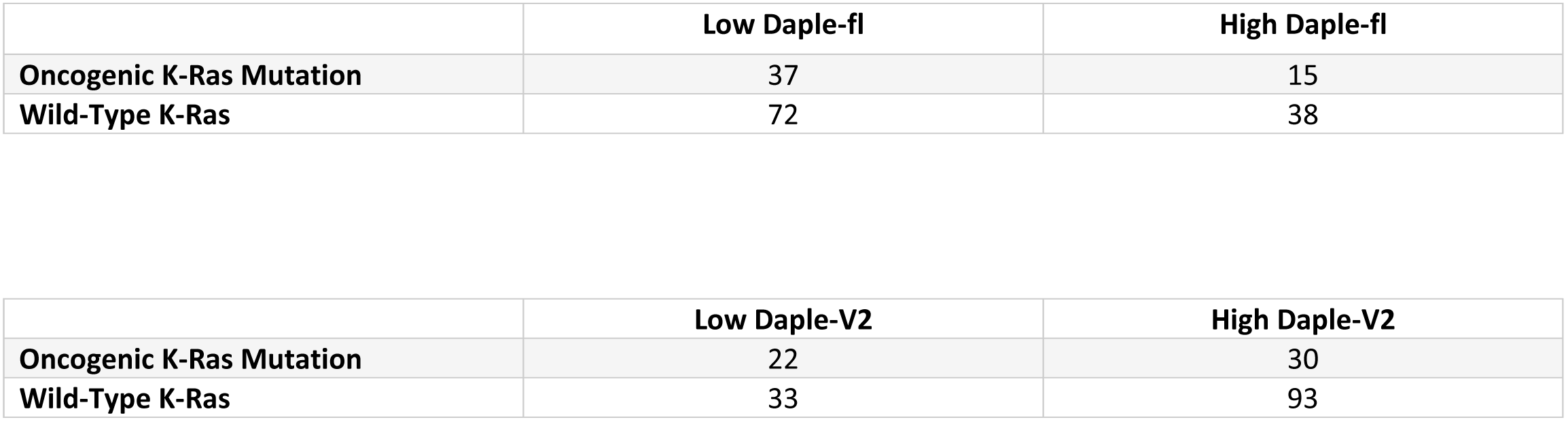
Contingency analysis (Fisher’s exact test) comparing Daple-fl or Daple-V2 expression and the presence of wild-type (WT) or oncogenic K-Ras mutation in 173 stage II colorectal carcinomas.

TABLES and LEGENDS
|  | Low Daple-fl | High Daple-fl |
| --- | --- | --- |
| **Oncogenic K-Ras Mutation** | 37 | 15 |
| **Wild-Type K-Ras** | 72 | 38 |

|  | Low Daple-V2 | High Daple-V2 |
| --- | --- | --- |
| **Oncogenic K-Ras Mutation** | 22 | 30 |
| **Wild-Type K-Ras** | 33 | 93 |

**Table 2.** Pearson’s correlation comparing Daple-fl or Daple-V2 expression and several tumor markers or histopathological parameters in 173 stage II colorectal carcinomas. Significant correlations are highlighted in red font. Daple-V2 showed significant negative correlation with Ki76 mitotic index and significant positive correlations with tumor suppressor SASH1 and serum levels of carcinoembryonic antigen (CEA).

|  | Ki67 Index | Osteopontin | SASH1 | MACC1 | Age | Tumor length (cm) | Tumor differentiation | Grading |
| --- | --- | --- | --- | --- | --- | --- | --- | --- |
| **Daple-V2** |  |  |  |  |  |  |  |  |
| **r-value** | -0.1794 | -0.01889 | 0.2041 | -0.1462 | 0.05122 | 0.01046 | -0.04673 | 0.02109 |
| **P value (two-tailed)** | **0.0245** | 0.8063 | **0.0074** | 0.0564 | 0.5059 | 0.892 | 0.5439 | 0.7842 |
| **Daple-fl** |  |  |  |  |  |  |  |  |
| **r-value** | 0.04645 | -0.03408 | 0.1224 | -0.1131 | 0.02617 | 0.007986 | -0.06434 | -0.00821 |
| **P value (two-tailed)** | 0.561 | 0.6553 | 0.1077 | 0.1373 | 0.7325 | 0.917 | 0.4004 | 0.9147 |

Finally, we previously discovered that high levels of expression of Daple in CTCs of patients with metastatic (Duke’s Stage IV) CRCs is associated with poorer prognosis compared to those with low Daple in CTCs (<u>Barbazan et al., 2016</u>); high Daple is associated with higher tumor recurrence at distant sites and poorer survival. Here we asked how each isoform contributed to the prognostic impact using the same cohort. We found that Daple-fl, but not Daple-V2, expression is increased in disseminated tumor cells compare to healthy subjects **(Fig 3-figure supplement 1A-B).** When we assessed their prognostic impact on survival, we found that although high expression of each isoform correlated with worse PFS **(Fig 3-figure supplement 1C, F)**, only Daple-fl levels carried a prognostic impact for DSS and overall survival (OS) **(Fig 3-figure supplement 1D-E, G-H)**. These findings are in keeping with our prior observations that pro-invasive and pro-EMT signatures are triggered exclusively by Daple-fl, but not Daple-V2 (**Fig 2M-Q**).

From these results we deduce that both Daple-fl and Daple-V2 cooperatively suppress tumor cell proliferation, that both transcripts are reduced during adenoma-to-carcinoma progression, and that the two isoforms have an additive prognostic impact, in that their reduced expression in tumors carries a poor prognosis. However, the two isoforms differ in their ability to trigger EMT and invasion; Daple-fl, but not Daple-V2 can support that.

### Daple is downregulated during adenoma-to-carcinoma conversion; the chemopreventive drug, Aspirin, has a differential effect on each isoform

Next we asked if suppression of Daple-fl and Daple-V2 transcripts during adenoma-to-cancer progression is associated also with reduced Daple proteins. Using an antibody raised against the C-terminus of Daple, which is identical between Daple-fl and Daple-V2, we confirmed that total Daple is expressed in the normal colon and in early and intermediate adenomas, but it is suppressed in advanced adenomas with villous features or high-grade dysplasia **(Fig 4A-B)**. In cancers, ~60% expressed Daple, but ~40% did not.

**Figure 4:**
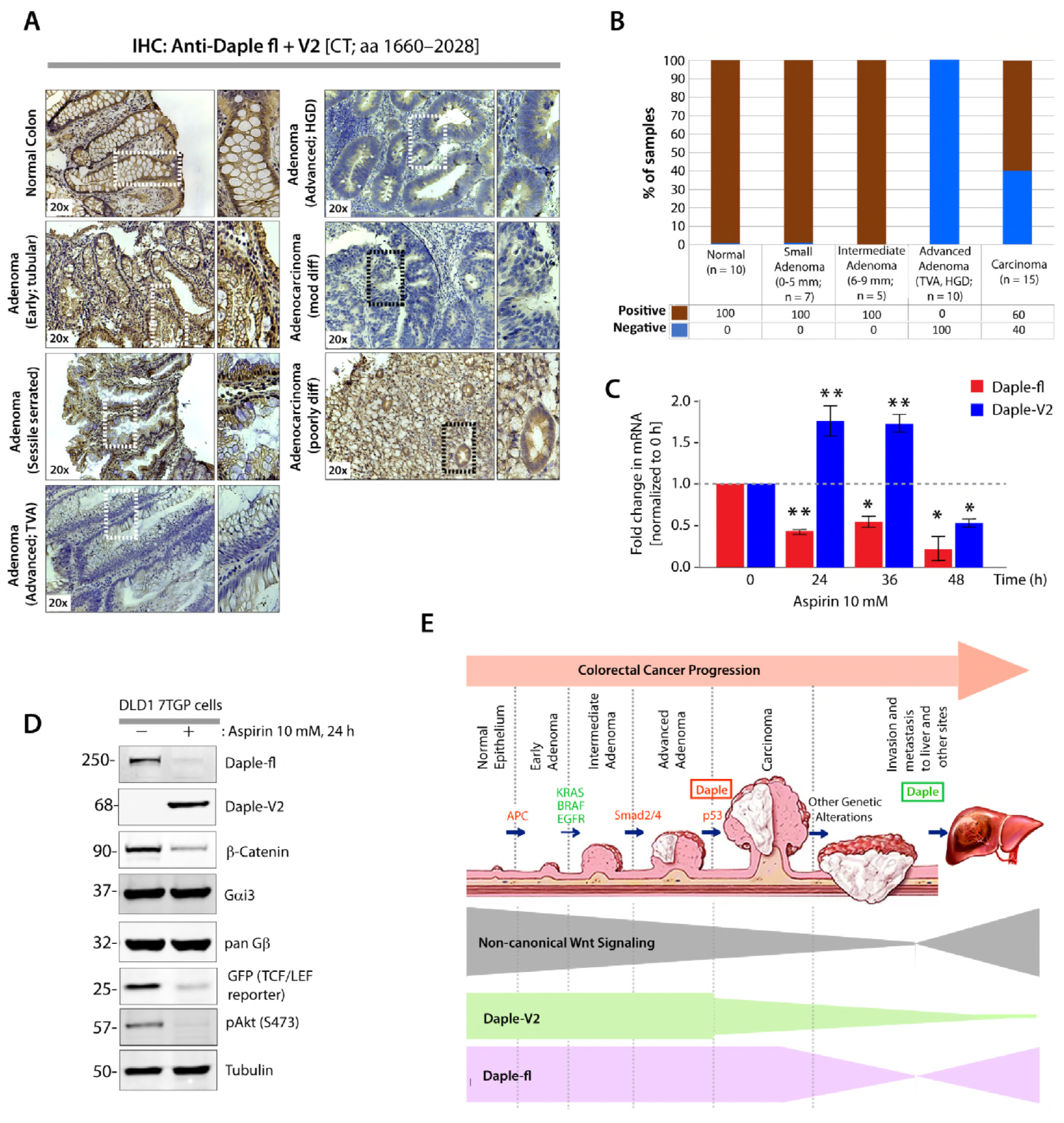
Both Daple-fl and Daple-V2 isoforms are downregulated during adenoma-to-carcinoma conversion and their expression is differentially regulated by the chemopreventive drug Aspirin. **A-B.** Expression of Daple protein was analyzed in formalin-fixed paraffin embedded human tissues (normal, adenomas and carcinomas) by immunohistochemistry (IHC) using anti-Daple-CT antibody that can detect both Daple-fl and Daple-V2 isoforms. *<u>Left</u>*: Representative tissues from each stained category are shown. Brown = positive stain. *<u>Right</u>*: Bar graphs display the proportion of samples in each category that stained positive vs. negative. **C.** Parental DLD1 cells were analyzed for Daple-fl and Daple-V2 mRNA by qPCR at indicated time Aspirin. Bar graphs display the fold change in each mRNA (Y axis) normalized to the expression levels at 0h. Error bars represent mean ± S.E.M of 3 independent experiments. *p* values: * = <0.05; ** < 0.01. **D.** Immunoblots showing the impact of low dose Aspirin on endogenous Daple isoforms expressed by DLD1-7TGP cells. β-Catenin and GFP were assessed as positive controls; the abundance of GFP serves as a surrogate marker for the transcriptional activity of β-Catenin via the TCF/LEF axis. G proteins, Gα and β-subunits were assessed as negative controls. **E**. Schematic summarizing profile of expression of Daple-fl and Daple-V2 isoforms during cancer initiation and metastatic progression in the colon. *<u>Upper</u>*: Various steps and histopathological stages of colorectal cancer progression are shown. Major genetic mutations/ deletions of key genes that herald the step-wise progression are indicated. Daple (both Daple-fl and Daple-V2) are decreased during adenoma to carcinoma progression (red box). Later, during cancer progression and systemic dissemination, total levels of Daple go up (green box), largely owing to an upregulation of its full-length (Daple-fl) transcript. *<u>Lower</u>*: Changes in the profile of expression of both Daple isoforms (Daple V2 = green; Daple-fl = purple) and their relationship to the previously identified patterns of non-canonical Wnt signaling (gray) is shown. CRC progression image courtesy of Johns Hopkins Digestive Disorders Library.

Next we asked if levels of expression of Daple-fl and Daple-V2 change in response to Aspirin, a potent chemopreventive agent that cuts the risk of CRCs by half (<u>Thun et al., 1991</u>; <u>Wunsch, 1998</u>). We found that treatment of DLD1 cells with low-dose Aspirin reduces both mRNA (**Fig 4C**) and protein (**Fig 4D**) for Daple-fl, which has both tumor-suppressive and pro-metastatic properties. By contrast, Aspirin increased the mRNA and protein for Daple-V2 (**Fig 4C-D**), which has only tumor-suppressive properties. Whether these observed changes in Daple isoforms are due to Aspirin’s ability to inhibit cyclo-oxygenase-II (COX2), i.e., COX2-dependent or independent mechanisms (<u>Goel et al., 2003</u>; <u>Wunsch, 1998</u>) remain unknown. Regardless, what is clear is that these changes are consistent with Aspirin’s ability to suppress polyp-to-cancer progression in the colon (<u>Barry et al., 2009</u>; <u>Johnson et al., 2010</u>), as well as its ability to inhibit metastatic progression of advanced CRCs (<u>Elwood et al., 2016</u>; <u>Guillem-Llobat et al., 2016</u>; <u>Jones et al., 1999</u>).

Taken together, our findings support the following model (**Fig 4E**). While both Daple-fl and Daple-V2 isoforms serve as growth suppressors early during oncogenesis, only Daple-fl serves as a pro-metastatic protein later during cancer progression. While both are suppressed during the step of polyp-to-cancer conversion, only Daple-fl is induced later during cancer invasion and dissemination.

## Conclusions

The major discovery we report here is the identification and characterization of a novel physiologic isoform of Daple, Daple-V2, which appears to contain the minimal modules that enable Daple to antagonize canonical β-Catenin-dependent Wnt signals and inhibit tumor cell growth and proliferation. Both isoforms are reduced during polyp-to-cancer progression in the colon. Compared to Daple-fl, Daple-V2 appears to be a more potent tumor suppressor and a better prognostic marker in primary tumors.

What Daple-V2 lacks is the ability to trigger EMT and invasion, which is a feature unique to Daple-fl. Consistent with these findings, we also found that although both isoforms collaboratively suppress tumor cell growth, and have an additive prognostic impact, only Daple-fl is increased in invasive margins and in CTCs disseminated during metastatic progression. Because we previously showed that a functional GBA motif is essential for Daple-fl to trigger EMT and invasion (<u>Aznar et al., 2015a</u>), findings we report here suggests that these functions require the coupling of Daple’s C-terminal G protein regulatory functions to the functions of its N-terminal HOOK and coiled-coil domains, e.g., phosphoinositide (PI3P) binding and localization to the pericentriolar recycling endosomes (<u>Aznar et al., 2017</u>), or binding to the dynein-dynactin motor complex (<u>Redwine et al., 2017</u>).

Although it remains unknown how cells modulate the expression of each Daple isoform, our studies provided precious clues; cells responding to Aspirin suppressed Daple-fl transcript and protein but concomitantly increased Daple-V2. Based on the properties of each isoform, we conclude that such a tradeoff is expected to overall maximize the desirable tumor-suppressive effect of Daple without the undesirable pro-EMT and pro-invasive effects.

In conclusion, we have revealed one of perhaps many ways how expression of Daple and its functions are regulated in the colon. Our findings that Daple isoforms can be pharmacologically manipulated to selectively augment its tumor-suppressive functions, while suppressing its pro-metastatic functions raises hope that therapeutic strategies could be tailored to meet such goals.

## ACKNOWLEDGMENTS

This work was supported by NIH grants CA100768, CA160911 and DK099226 (to P.G). P.G. was also supported by the American Cancer Society (ACS-IRG 70-002) and by the UC San Diego Moores Cancer Center. J.E was supported by a NCI/NIH-funded Cancer Biology, Informatics & Omics (CBIO) Training Program (T32 CA067754) and a Postdoctoral Fellowship, PF-18-101-01-CSM, from the American Cancer Society, Y.M was supported by the NIH Training Grant in Gastroenterology (T32DK0070202) and a Clinical and Translational Science Awards (CTSA) Program (TL1TR001443), and J.B. by a fellowship from the Spanish Ministry of Education, Culture and Sports (FPU, AP2009-5229). Other sources of funding include NIH R01CA72851 (to A.G), the German ministry of education and research (BMBF/m4 Biobank Alliance) and funds from the Kommission für klinische Forschung (to U.N and K.P-J).

## AUTHOR CONTRIBUTIONS

Y.D, N.A., J.E. and P.G designed, performed and analyzed most of the experiments in this work. N.A and Y.D carried out the biochemical and cell biological characterization studies comparing and contrasting the two Daple isoforms. Y.D., Y.M. and P.G carried out the IHC studies on FFPE patient tissues. U.N. and K-P.J. provided access to CRC Stage-II cohort and primary tumor-derived RNA and U.N. and Y.D generated and analyzed the data. J.B. and Y.D carried out the analyses of Daple in CTCs. A.G. provided access to adenomas, and contributed unpublished essential data or reagents. Y.D., N.A and P.G conceived the project and wrote the manuscript. P.G supervised and funded the project.

## Materials and Methods

### Reagents and Antibodies

Unless otherwise indicated, all reagents were of analytical grade and obtained from Sigma-Aldrich. Cell culture media were purchased from Invitrogen. All restriction endonucleases and Escherichia coli strain DH5α were purchased from New England Biolabs. E. coli strain BL21 (DE3), phalloidin-Texas Red were purchased from Invitrogen. Genejuice transfection reagent was from Novagen. PfuUltra DNA polymerase was purchased from Stratagene. Recombinant Goat anti-rabbit and goat anti-mouse Alexa Fluor 680 or IRDye 800 F(ab′)2 used for immunoblotting were from Li-Cor Biosciences. Mouse anti-αtubulin and anti-actin were obtained from Sigma; anti-Myc was obtained from Covance, and anti-GFP from Santa Cruz Biotechnology. Rabbit anti–pan-Gβ (M-14), and anti-Gαi3 were obtained from Santa Cruz Biotechnology. Anti-Daple antibodies were generated in collaboration with Millipore using the C-terminus of Daple (aa 1660-2028) as an immunogen.

### Plasmid Constructs and Mutagenesis

Cloning of N-terminally tagged myc-Daple-fl in pcDNA3.1(+) was carried out as described previously (<u>Aznar et al., 2015b</u>). The subsequent site-directed mutagenesis and truncated constructs (myc-Daple full length F1675A (FA) and deleted from aa 2025-2028(ΔPBM) were carried out on this template using Quick Change as per manufacturer’s protocol. Cloning of N-terminally tagged myc-Daple-V2 was carried out by PCR cloning directly from myc-pcDNA3.1(+)-Daple-fl using primers containing 5’ unique region ( MSVLS) in Daple V2 sequence ( corresponding to UniProtKB - Q9P219-2) and being inserted into myc-pcDNA 3.1 (+) between Kpn-1/EcoR1. Daple-V2-FA (F169A) and Daple-V2- ΔPBM (deleted from 549- 552aa) were alos directly cloned out from pcDNA3.1(+)-Daple-fl-FA and pcDNA3.1(+)-daple-fl-ΔPBM. . Cloning of rat Gα-proteins into pGEX-4T-1 GST-Gαi3 has been described previously(<u>Garcia-Marcos et al., 2010</u>; <u>Garcia-Marcos et al., 2009</u>; <u>Garcia-Marcos et al., 2011b</u>; <u>Ghosh et al., 2010</u>; <u>Ghosh et al., 2008</u>). GST-tagged FZDR7-CT construct (<u>Yao et al., 2004</u>) was a generous gift from Ryoji Yao (JFCR research institute, Japan). GST-Dvl2-PDZ was from Raymond Habas (Temple University, Philadelphia, PA).

### Protein Expression and Purification

GST, GST-Gαi3, GST-PDZ and GST-FZDR7 fusion constructs were expressed in *E. coli* strain BL21 (DE3) (Invitrogen) and purified as described previously (<u>Garcia-Marcos et al., 2009</u>; <u>Ghosh et al., 2010</u>; <u>Ghosh et al., 2008</u>). Briefly, bacterial cultures were induced overnight at 25°C with 1 mM isopropyl β-D-1-thiogalactopyranoside (IPTG). Pelleted bacteria from 1L of culture were re-suspended in 10 ml GST-lysis buffer [25 mMTris-HCl, pH 7.5, 20 mMNaCl, 1 mM EDTA, 20% (v:v) glycerol, 1% (v:v) Triton X-100, 2X protease inhibitor cocktail (Complete EDTA-free, Roche Diagnostics). After sonication (4 x 20s, 1 min between cycles), lysates were centrifuged at 12,000g at 4°C for 20 min. Except for GST-FZD (see *in vitro* GST pulldown assay section), solubilized proteins were affinity purified on glutathione-Sepharose 4B beads (GE Healthcare). Proteins were eluted, dialyzed overnight against PBS and stored at -80 °C.

### Cell Culture and the Rationale for Choice of Cells in Various Assays

Tissue culture was carried out essentially as described before (<u>Garcia-Marcos et al., 2011a</u>; <u>Ghosh et al., 2010</u>; <u>Ghosh et al., 2008</u>). We used a total of 3 different cell lines in this work, each chosen carefully based on its level of endogenous Daple expression and the type of assay. All these cell lines were cultured according to ATCC guidelines. Cos7 cells were primarily used for transient overexpression of tagged Daple protein and lysates of these cells were used as source of proteins in pulldown assays. We chose to carry out these assays in Cos7 cells because they are easily and efficiently transfected (> 90% efficiency) with most constructs. The added advantage is that they have no detectable endogenous Daple and provide a system to selectively analyze the properties of WT vs mutant Daple constructs without interference from endogenous Daple.

DLD1 were primarily used to study the effect of Daple on cancer cell growth properties (anchorage-dependent) and to assess the effect of Daple on the classical Wnt signaling pathway (βCatenin/TCF/LEF). There are several reasons why this cell line was chosen: 1) We focused on colorectal cancer in this study and DLD1 cells were appropriate to translate our findings because they are human colorectal cancer cells; 2) We determined that levels of Daple are significantly lower/undetectable (~ 10 fold) in these cells compared to normal colon (data not shown), thereby allowing us to reconstitute Daple expression exogenously and analyze the effect of various mutant Daple constructs without significant interference due to the endogenous protein; 3) These cells have been extensively characterized with respect to most oncogenes (ATCC database), and are highly tumorigenic in 2-D and 3-D cultures due to a mutation in KRAS (G13D) (Shirasawa et al., 1993; Ahmedet al., 2013); 4) They are a sensitive model to study how various manipulations of the noncanonical Wnt signaling pathway oppose the canonical Wnt pathway during tumor growth because they constitutively secrete Wnt ligands to maintain high levels of the canonical signaling(<u>Voloshanenko et al., 2013</u>) within the growth matrix. Production and secretion of endogenous ligands bypasses the need to add exogenous ligands repeatedly during prolonged assays that last ~2 weeks.

Low passage NIH3T3 fibroblasts were used exclusively in 3-D Matrigel invasion. The rationale for their use in invasion assay lies in the fact that non-transformed NIH3T3 fibroblasts are poorly invasive in vitro and non-tumorigenic and non-metastatic in animal studies (<u>Bondy et al., 1985</u>; <u>Chambers et al., 1990</u>; <u>Hill et al., 1988</u>; <u>Tuck et al., 1991</u>). It is because of this reason, NIH3T3 cells are widely used to study proteins that can trigger a gain in invasive properties (<u>Leitner et al., 2011</u>). The rationale for using NIH3T3 in the above assays is further strengthened by the fact that they are highly transfectable (~80% transfection efficiency with myc-Daple) and express Daple at very low endogenous levels (as determined by immunoblotting and qPCR) compared to normal colonic epithelium. Such expression pattern allows us to study the effect of various mutant Daple constructs without significant interference due to the endogenous protein.

### Transfection; Generation of Stable Cell Lines and Cell Lysis

Transfection was carried out using Genejuice (Novagen) for DNA plasmids following the manufacturers’ protocols. DLD1 cell lines stably expressing Daple constructs were selected after transfection in the presence of 800 μg/ml G418 for 6 weeks. The resultant multiclonal pool was subsequently maintained in the presence of 500 μg/ ml G418. Daple expression was verified independently using anti-Daple antibody by immunoblotting, and estimated to be ~5x the endogenous level.

### Quantitative Immunoblotting

For immunoblotting, protein samples were separated by SDS-PAGE and transferred to PVDF membranes (Millipore). Membranes were blocked with PBS supplemented with 5% non fat milk (or with 5% BSA when probing for phosphorylated proteins) before incubation with primary antibodies.Infrared imaging with two-color detection and band densitometry quantifications were performed using a Li-Cor Odyssey imaging system exactly as done previously (<u>Garcia-Marcos et al., 2011a</u>; <u>Garcia-Marcos et al., 2010</u>; <u>Garcia-Marcos et al., 2012</u>; <u>Garcia-Marcos et al., 2011b</u>; <u>Ghosh et al., 2010</u>). All Odyssey images were processed using Image J software (NIH) and assembled into figure panels using Photoshop and Illustrator software (Adobe).

### In vitro GST pulldown

Purified GST alone, GST-Gαi3 or GST-PDZ (5 μg) were immobilized on glutathione-Sepharose beads and incubated with binding buffer [50 mM Tris-HCl (pH 7.4), 100 mM NaCl, 0.4% (v:v) Nonidet P-40, 10 mM MgCl_2_, 5 mM EDTA, 30 μM GDP, 2 mM DTT, protease inhibitor mixture] for 90 min at room temperature as described before (<u>Garcia-Marcos et al., 2011a</u>; <u>Ghosh et al., 2010</u>; <u>Ghosh et al., 2008</u>; <u>Lin et al., 2011</u>). Lysates (~250 μg) of Cos7 cells expressing appropriate myc-Daple-V2 constructs were added to each tube, and binding reactions were carried out for 4 h at 4°C with constant tumbling in binding buffer [50 mM Tris-HCl (pH 7.4), 100 mM NaCl, 0.4% (v:v) Nonidet P-40, 10 mM MgCl_2_, 5 mM EDTA, 30 μM GDP, 2 mM DTT]. Beads were washed (4x) with 1 mL of wash buffer [4.3 mM Na_2_HPO_4_, 1.4 mM KH_2_PO_4_ (pH 7.4), 137 mM NaCl, 2.7 mM KCl, 0.1% (v:v) Tween 20, 10 mM MgCl_2_, 5 mM EDTA, 30 μM GDP, 2 mM DTT] and boiled in Laemmli’s sample buffer. Immunoblot quantification was performed by infrared imaging following the manufacturer’s protocols using an Odyssey imaging system (Li-Cor Biosciences).

GST-FZD7-CT construct was immobilized on glutathione-Sepharose beads directly from bacterial lysates by overnight incubation at 4°C with constant tumbling as described before (<u>Aznar et al., 2015b</u>). Next morning, GST-FZD7-CT immobilized on glutathione beads were washed and subsequently incubated with His-tagged Daple-CT or Gαi3 proteins at 4°C with constant tumbling. Washes and immunoblotting were performed as previously.

### β-Catenin Reporter Assays

These assays were carried out using the well-established reporter 7xTcf-eGFP(7TGP) (<u>Fuerer and Nusse, 2010</u>). Stable cells lines expressing this reporter were generated by lentiviral transduction and subsequent selection using standard procedures. Lentiviral infection and selection were performed according to standard procedures. Briefly, 10 cm plates DLD1 cells at 70% confluency were incubated with media containing 8 μg/mL polybrene and 10 μl of lentivirus for 6 h. After 24 hours post infection, selection of puromycin-resistant clones was initiated by adding the antibiotic at 2 μg/ml final concentration. The resultant DLD1-7TGP stable cells were subsequently transfected with various myc-Daple V2 constructs and selected for G418 resistance as described earlier in methods. The DLD1-7TGP cells stably expressing myc-Daple were incubated overnight at 0.2% FBS, analyzed by fluorescence microscopy, and photographed prior to lysis. Whole cell lysates samples were then boiled in Laemmli’s sample buffer and GFP protein expression was monitored by immunoblotting..

### Anchorage-dependent Colony Growth Assay

Anchorage-dependent growth was monitored on solid (plastic) surface. Approximately ~1000 DLD1 cells stably expressing various Daple constructs were plated in 6-well plates and incubated in 5% CO_2_ at 37°C for ~2 weeks in 0.2% FBS growth media. Colonies were then stained with 0.005% crystal violet for 1 h. The remaining DLD1 cells were lysed and analyzed by WB to confirm Daple construct expression. Each experiment was analyzed in triplicate.

### Invasion Assays

NIH3T3 cell invasion assay in 3D culture was performed according to the manufacturer’s protocol (Trevigen, Cultrex 3D Spheroid BME Cell Invasion Assay, catalog # 3500-096-K). Briefly, NIH3T3 cells (3000 cells) transfected with empty vector (control) or myc-Daple constructs were incubated first in the Spheroid Formation extracellular matrix (ECM) containing 0.2% FBS for 3 days. Invasion matrix was then added and layered on top with media containing FBS. Serum-triggered cell invasion was photographed under light microscope everyday for 10 days and fresh media (FBS concentration is increased each time in order to maintain a gradient) was replenished every 48 h. Photographs were analyzed and pseudocolored by Image J to reflect cell density.

### Patient cohort for mRNA analysis

The ethics committee of the Klinikumrechts der Isar, Munich, Germany, approved collection of the patient samples (#1926/07, and #5428/12). All samples were obtained after prior informed written consent. Tumor tissue from 173 patients with histopathologically confirmed stage II (AJCC/UICC) colon cancer who underwent complete surgical resection (R0) between 1987 and 2006 was obtained, by a pathologist immediately after surgical resection. Specimens were transferred into liquid nitrogen and stored at -80°C until further processing. None of the patients received neoadjuvant treatment. No metachronous tumors were found in the colon or rectum. As reported previously in detail, clinical data and post-operative follow-up was collected for all patients; moreover, DNA was isolated for KRAS and BRAF mutation analysis, as well as microsatellite instability testing (<u>Nitsche et al., 2012</u>). Total mRNA was extracted by standard procedures (Qiagen, Hilden, Germany), after histology guided sample selection to ensure a tumor cell content of >50%, and transcribed to cDNA as described in detail earlier, for expression analysis of DAPLE (<u>Nitsche et al., 2012</u>).

### RNA isolation, standard curve and quantitative PCR (qPCR)

Total RNA was isolated using an RNeasy kit (QIAGEN) as per the manufacturers’ protocol. First-strand cDNA was synthesized using Superscript II reverse transcriptase (Invitrogen), followed by ribonuclease H treatment (Invitrogen) prior to performing quantitative real-time PCR. A standard curve, to quantify mRNA copy number, was constructed using larger PCR products (~700bp) that included the target sequence used in qPCR. Reactions omitting reverse transcriptase were performed in each experiment as negative controls. Reactions were then run on a real-time PCR system (ABI StepOnePlus; Applied Biosystems). Gene expression was detected with SYBR green/Taqman assay (Invitrogen), and relative gene expression was determined by normalizing to GAPDH using the comparative ΔCt/Relative standard curve method.

Primer and probe sequences are as listed below.

**Probes and primers used in Taqman assays:** for human tumor/tissue and CTC samples

GAPDH:

hGAPDH-fwd: 5’- CAGTTGTAGGCAAGCTGCGA -3’ hGAPDH-rev: 5’- TATGACAGGCCCGAAGCTTCT -3’ hGAPDH-probe: 5’- CCAAGCCTGAGGGCAAGGCTATAATAGATGAAT-3’ hGAPDH-standard fwd: 5’- GCT GTG ACA TCA GGG CAA T-3’ hGAPDH-standard rev: 5’ - GGC GGT GGT GGC TTT ATT T-3’

Daple-fl:

hDaple-fl-fwd: 5’- CGGGACCTCACCAAGCAA -3’ hDaple-fl-rev: 5’- CTGCTGAGCTGCTGGCTCTT -3’ hDaple-fl - probe: 5’- CAACTCTGAGGGAGGACCTGGTGCTC -3’ hDaple-fl-Standard-fwd: 5’- GGATGCAGTCTTGGACGATAG -3’ hDaple-fl-Standard-rev: 5’- CTTCTTTCATGGCTAGTGTTGTTT -3’

Daple-V2:

hDaple-V2-fwd: 5’- GGAGCCTCAGGATATACGTGCA -3’ hDaple-V2-rev: 5’- TCAAGGCTGCCTCTGTGTGG -3’ hDaple-V2-probe: 5’- CAGGATGTCCGTACTAAGCCCTGGGGATC -3’ hDaple-V2 Standard-fwd: 5’- CACTCCCTGGACCATTTCTT -3’ hDaple-V2 Standard-rev: 5’- CTGTAGTGGTGGCTGAAGTT -3’

**Primers used in SYBR-green assays:** for cell-based analyses

Lox-3:

hq-LOXL3 fwd: 5’- ATGGGTGCTATCCACCTGAG -3’ hq-LOXL3 rev: 5’- GAGTCGGATCCTGGTCTCTG –3’

Axin-2:

hAxin-2-fwd: 5’- GAGTGGACTTGTGCCGACTTCA -3’ hAxin-2-rev: 5’- GGTGGCTGGTGCAAAGACATAG -3’

Vimentin:

hVim-fwd: 5’- AAGAGAACTTTGCCGTTGAA-3’ hVim-rev: 5’-GTGATGCTGAGAAGTTTCGT-3’

SFRP-1:

hSFRP-1-fwd: 5’- GAGTTTGCACTGAGGATGAAAA -3’ hSFRP-1-rev: 5’- GCTTCTTCTTCTTGGGGACA -3’

GADPH:

hGADPH q-fwd2: 5’- TCA GTT GTA GGC AAG CTG CGA CGT-3’ hGADPH q-rev2: 5’- AAGCCAGAGGCTGGTACCTAGAAC-3

DAPLE:

hDaple-fl-fwd: 5’- TGA CAT GGA GAC CCT GAA GGC TGA -3’ hDaple fl-rev2: 5’- TTTCATGCGGGCCTCACTGCTGA-3’ hDaple V2-Fwd : 5’- GTT GTC ACA CTC CCT GGA CCA TTT C -3’ hDaple V2-Rev: 5’- GCTTTGGTTTTAGATCCCCAGGGC -3’

### Patient cohort for IHC analysis

Formalin-fixed paraffin embedded (FFPE) normal, polyp and cancer tissues used in this study were obtained from patients undergoing routine colonoscopies and provided by the section of Gastroenterology, VA San Diego Healthcare System. The protocol was approved by the Human Research Protection Program Institutional Review Board (protocol H130266).

### Immunohistochemistry

Colon specimens of known histologic type were analyzed by IHC using a previously-validated anti-Daple-CT rabbit polyclonal antibody raised against Daple (aa 1660-2028) (1:50; Millipore Inc.). Briefly, formalin-fixed, paraffin-embedded tissue sections of 4 μm thickness were cut and placed on glass slides coated with 3-aminopropyl triethoxysilane, followed by deparaffinization and hydration. Heat-induced epitope retrieval was performed using citrate buffer (pH 6) in a pressure cooker. Tissue sections were incubated with 3% hydrogen peroxidase for 15 min to block endogenous peroxidase activity, followed by incubation with primary antibodies overnight in a humidified chamber at 4°C. Immunostaining was visualized with a labeled streptavidin-biotin using 3,3′-diaminobenzidine as a chromogen and counterstained with hematoxylin. All the samples were first quantitatively analyzed and scored based on 2 independent criteria. First, the intensity of staining was scored on a scale of 0 to 3, where 0 = no staining, 1 = light brown, 2 = brown, and 3 = dark brown. Second, the percentage of the cells that stained positive in the tumor area was scored on a scale of 0 to 4, where 0 = 0, 1 = ≤10%, 2 = 11–50%, 3 = 51–75%, and 4 = >75%. Subsequently, each tumor sample was assigned a final score, which is the product of its (intensity of staining) × (% cells that stained positive). Tumors were categorized as negative when their final score was <3 and as positive when their final score was ≥3.

### Statistical analyses

Statistical evaluation was performed using GraphPad Prism 5 software. Unless stated otherwise, statistical significance was determined using Student’s *t* test. The associations between the expression level of Daple isoforms and K-ras mutation status and metastatic status of disease were investigated by Fisher’s exact test. Parametric Pearson’s correlation and nonparametric Spearman’s correlation analysis were used to assess the relationship between the expression of Daple isoforms and patients’ age, tumor size, differentiation and grading, the level of Ki67-Index, Osteopontin, SASH1, MACC1 and CEA. In order to derive optimal cut-off values of gene expression levels, maximally selected log-rank statistics performed by R Software version 2.15.0.(<u>Budczies et al., 2012</u>) were used. Time-dependent survival probabilities were estimated with the Kaplan-Meier method using the log-rank test. All statistical tests were performed two-sided, and p-values less than 0.05 were considered to be statistically significant.

## LITERATURE CITED

Aznar, N., J. Ear, Y. Dunkel, N. Sun, K. Satterfield, F. He, N.A. Kalogriopoulos, I. Lopez-Sanchez, M. Ghassemian, D. Sahoo, I. Kufareva, and P. Ghosh. 2018. Convergence of Wnt, growth factor, and heterotrimeric G protein signals on the guanine nucleotide exchange factor Daple. Sci Signal. 11.

Aznar, N., K.K. Midde, Y. Dunkel, I. Lopez-Sanchez, Y. Pavlova, A. Marivin, J. Barbazan, F. Murray, U. Nitsche, K.P. Janssen, K. Willert, A. Goel, M. Abal, M. Garcia-Marcos, and P. Ghosh. 2015a. Daple is a novel non-receptor GEF required for trimeric G protein activation in Wnt signaling. Elife. 4:e07091.

Aznar, N., K.K. Midde, Y. Dunkel, I. Lopez-Sanchez, Y. Pavlova, A. Marivin, J. Barbazan, F. Murray, U. Nitsche, K.P. Janssen, K. Willert, A. Goel, M. Abal, M. Garcia-Marcos, and P. Ghosh. 2015b. Daple is a novel non-receptor GEF required for trimeric G protein activation in Wnt signaling. eLife. 4.

Aznar, N., N. Sun, Y. Dunkel, J. Ear, M.D. Buschman, and P. Ghosh. 2017. A Daple-Akt feed-forward loop enhances noncanonical Wnt signals by compartmentalizing beta-catenin. Mol Biol Cell. 28:3709–3723.

Barbazan, J., Y. Dunkel, H. Li, U. Nitsche, K.P. Janssen, K. Messer, and P. Ghosh. 2016. Prognostic Impact of Modulators of G proteins in Circulating Tumor Cells from Patients with Metastatic Colorectal Cancer. Sci Rep. 6:22112.

Barry, E.L., L.B. Sansbury, M.V. Grau, I.U. Ali, S. Tsang, D.J. Munroe, D.J. Ahnen, R.S. Sandler, F. Saibil, J. Gui, R.S. Bresalier, G.E. McKeown-Eyssen, C. Burke, and J.A. Baron. 2009. Cyclooxygenase-2 polymorphisms, aspirin treatment, and risk for colorectal adenoma recurrence--data from a randomized clinical trial. Cancer Epidemiol Biomarkers Prev. 18:2726–2733.

Bondy, G.P., S. Wilson, and A.F. Chambers. 1985. Experimental metastatic ability of H-ras-transformed NIH3T3 cells. Cancer research. 45:6005–6009.

Budczies, J., F. Klauschen, B.V. Sinn, B. Gyorffy, W.D. Schmitt, S. Darb-Esfahani, and C. Denkert. 2012. Cutoff Finder: a comprehensive and straightforward Web application enabling rapid biomarker cutoff optimization. PloS one. 7:e51862.

Chambers, A.F., G.H. Denhardt, and S.M. Wilson. 1990. ras-transformed NIH 3T3 cell lines, selected for metastatic ability in chick embryos, have increased proportions of p21-expressing cells and are metastatic in nude mice. Invasion & metastasis. 10:225–240.

Ellis, M.J., V.J. Suman, J. Hoog, R. Goncalves, S. Sanati, C.J. Creighton, K. DeSchryver, E. Crouch, A. Brink, M. Watson, J. Luo, Y. Tao, M. Barnes, M. Dowsett, G.T. Budd, E. Winer, P. Silverman, L. Esserman, L. Carey, C.X. Ma, G. Unzeitig, T. Pluard, P. Whitworth, G. Babiera, J.M. Guenther, Z. Dayao, D. Ota, M. Leitch, J.A. Olson, Jr., D.C. Allred, and K. Hunt. 2017. Ki67 Proliferation Index as a Tool for Chemotherapy Decisions During and After Neoadjuvant Aromatase Inhibitor Treatment of Breast Cancer: Results From the American College of Surgeons Oncology Group Z1031 Trial (Alliance). J Clin Oncol. 35:1061–1069.

Elwood, P.C., G. Morgan, J.E. Pickering, J. Galante, A.L. Weightman, D. Morris, M. Kelson, and S. Dolwani. 2016. Aspirin in the Treatment of Cancer: Reductions in Metastatic Spread and in Mortality: A Systematic Review and Meta-Analyses of Published Studies. PloS one. 11:e0152402.

Fuerer, C., and R. Nusse. 2010. Lentiviral vectors to probe and manipulate the Wnt signaling pathway. PloS one. 5:e9370.

Garcia-Marcos, M., J. Ear, M.G. Farquhar, and P. Ghosh. 2011a. A GDI (AGS3) and a GEF (GIV) regulate autophagy by balancing G protein activity and growth factor signals. Mol Biol Cell. 22:673–686.

Garcia-Marcos, M., P. Ghosh, J. Ear, and M.G. Farquhar. 2010. A structural determinant that renders G alpha(i) sensitive to activation by GIV/girdin is required to promote cell migration. J Biol Chem. 285:12765–12777.

Garcia-Marcos, M., P. Ghosh, and M.G. Farquhar. 2009. GIV is a nonreceptor GEF for G alpha i with a unique motif that regulates Akt signaling. Proc Natl Acad Sci U S A. 106:3178–3183.

Garcia-Marcos, M., P.S. Kietrsunthorn, Y. Pavlova, M.A. Adia, P. Ghosh, and M.G. Farquhar. 2012. Functional characterization of the guanine nucleotide exchange factor (GEF) motif of GIV protein reveals a threshold effect in signaling. Proc Natl Acad Sci U S A. 109:1961–1966.

Garcia-Marcos, M., P.S. Kietrsunthorn, H. Wang, P. Ghosh, and M.G. Farquhar. 2011b. G Protein binding sites on Calnuc (nucleobindin 1) and NUCB2 (nucleobindin 2) define a new class of G(alpha)i-regulatory motifs. J Biol Chem. 286:28138–28149.

Ghosh, P., A.O. Beas, S.J. Bornheimer, M. Garcia-Marcos, E.P. Forry, C. Johannson, J. Ear, B.H. Jung, B. Cabrera, J.M. Carethers, and M.G. Farquhar. 2010. A G{alpha}i-GIV molecular complex binds epidermal growth factor receptor and determines whether cells migrate or proliferate. Mol Biol Cell. 21:2338–2354.

Ghosh, P., M. Garcia-Marcos, S.J. Bornheimer, and M.G. Farquhar. 2008. Activation of Galphai3 triggers cell migration via regulation of GIV. J Cell Biol. 182:381–393.

Goel, A., D.K. Chang, L. Ricciardiello, C. Gasche, and C.R. Boland. 2003. A novel mechanism for aspirin-mediated growth inhibition of human colon cancer cells. Clin Cancer Res. 9:383–390.

Guillem-Llobat, P., M. Dovizio, A. Bruno, E. Ricciotti, V. Cufino, A. Sacco, R. Grande, S. Alberti, V. Arena, M. Cirillo, C. Patrono, G.A. FitzGerald, D. Steinhilber, A. Sgambato, and P. Patrignani. 2016. Aspirin prevents colorectal cancer metastasis in mice by splitting the crosstalk between platelets and tumor cells. Oncotarget. 7:32462–32477.

Hill, S.A., S. Wilson, and A.F. Chambers. 1988. Clonal heterogeneity, experimental metastatic ability, and p21 expression in H-ras-transformed NIH 3T3 cells. Journal of the National Cancer Institute. 80:484–490.

Ishida-Takagishi, M., A. Enomoto, N. Asai, K. Ushida, T. Watanabe, T. Hashimoto, T. Kato, L. Weng, S. Matsumoto, M. Asai, Y. Murakumo, K. Kaibuchi, A. Kikuchi, and M. Takahashi. 2012. The Dishevelled-associating protein Daple controls the non-canonical Wnt/Rac pathway and cell motility. Nature communications. 3:859.

Johnson, C.C., R.B. Hayes, R.E. Schoen, M.J. Gunter, W.Y. Huang, and P.T. Team. 2010. Non-steroidal anti-inflammatory drug use and colorectal polyps in the Prostate, Lung, Colorectal, And Ovarian Cancer Screening Trial. Am J Gastroenterol. 105:2646–2655.

Jones, M.K., H. Wang, B.M. Peskar, E. Levin, R.M. Itani, I.J. Sarfeh, and A.S. Tarnawski. 1999. Inhibition of angiogenesis by nonsteroidal anti-inflammatory drugs: insight into mechanisms and implications for cancer growth and ulcer healing. Nat Med. 5:1418–1423.

Kobayashi, H., T. Michiue, A. Yukita, H. Danno, K. Sakurai, A. Fukui, A. Kikuchi, and M. Asashima. 2005. Novel Daple-like protein positively regulates both the Wnt/beta-catenin pathway and the Wnt/JNK pathway in Xenopus. Mechanisms of development. 122:1138–1153.

Larochelle, S. 2016. SYSTEMS BIOLOGY. Protein isoforms: more than meets the eye. Nat Methods. 13:291.

Leitner, L., D. Shaposhnikov, A. Mengel, A. Descot, S. Julien, R. Hoffmann, and G. Posern. 2011. MAL/MRTF-A controls migration of non-invasive cells by upregulation of cytoskeleton-associated proteins. Journal of cell science. 124:4318–4331.

Lin, C., J. Ear, Y. Pavlova, Y. Mittal, I. Kufareva, M. Ghassemian, R. Abagyan, M. Garcia-Marcos, and P. Ghosh. 2011. Tyrosine phosphorylation of the Galpha-interacting protein GIV promotes activation of phosphoinositide 3-kinase during cell migration. Sci Signal. 4:ra64.

Martini, M., A. Gnann, D. Scheikl, B. Holzmann, and K.P. Janssen. 2011. The candidate tumor suppressor SASH1 interacts with the actin cytoskeleton and stimulates cell-matrix adhesion. Int J Biochem Cell Biol. 43:1630–1640.

Niavarani, A., T. Herold, Y. Reyal, M.C. Sauerland, T. Buchner, W. Hiddemann, S.K. Bohlander, P.J. Valk, and D. Bonnet. 2016. A 4-gene expression score associated with high levels of Wilms Tumor-1 (WT1) expression is an adverse prognostic factor in acute myeloid leukaemia. British journal of haematology. 172:401–411.

Niikura, N., T. Iwamoto, S. Masuda, N. Kumaki, T. Xiaoyan, M. Shirane, K. Mori, B. Tsuda, T. Okamura, Y. Saito, Y. Suzuki, and Y. Tokuda. 2012. Immunohistochemical Ki67 labeling index has similar proliferation predictive power to various gene signatures in breast cancer. Cancer Sci. 103:1508–1512.

Nitsche, U., R. Rosenberg, A. Balmert, T. Schuster, J. Slotta-Huspenina, P. Herrmann, F.G. Bader, H. Friess, P.M. Schlag, U. Stein, and K.P. Janssen. 2012. Integrative marker analysis allows risk assessment for metastasis in stage II colon cancer. Annals of surgery. 256:763–771; discussion 771.

Oshita, A., S. Kishida, H. Kobayashi, T. Michiue, T. Asahara, M. Asashima, and A. Kikuchi. 2003. Identification and characterization of a novel Dvl-binding protein that suppresses Wnt signalling pathway. Genes to cells : devoted to molecular & cellular mechanisms. 8:1005–1017.

Redwine, W.B., M.E. DeSantis, I. Hollyer, Z.M. Htet, P.T. Tran, S.K. Swanson, L. Florens, M.P. Washburn, and S.L. Reck-Peterson. 2017. The human cytoplasmic dynein interactome reveals novel activators of motility. Elife. 6.

Thun, M.J., M.M. Namboodiri, and C.W. Heath, Jr. 1991. Aspirin use and reduced risk of fatal colon cancer. N Engl J Med. 325:1593–1596.

Tuck, A.B., S.M. Wilson, R. Khokha, and A.F. Chambers. 1991. Different patterns of gene expression in ras-resistant and ras-sensitive cells. Journal of the National Cancer Institute. 83:485–491.

Voloshanenko, O., G. Erdmann, T.D. Dubash, I. Augustin, M. Metzig, G. Moffa, C. Hundsrucker, G. Kerr, T. Sandmann, B. Anchang, K. Demir, C. Boehm, S. Leible, C.R. Ball, H. Glimm, R. Spang, and M. Boutros. 2013. Wnt secretion is required to maintain high levels of Wnt activity in colon cancer cells. Nature communications. 4:2610.

Wunsch, H. 1998. COX provides missing link in mechanism of aspirin in colon cancer. Lancet. 351:1864.

Yao, R., Y. Natsume, and T. Noda. 2004. MAGI-3 is involved in the regulation of the JNK signaling pathway as a scaffold protein for frizzled and Ltap. Oncogene. 23:6023–6030.

Zeller, C., B. Hinzmann, S. Seitz, H. Prokoph, E. Burkhard-Goettges, J. Fischer, B. Jandrig, L.E. Schwarz, A. Rosenthal, and S. Scherneck. 2003. SASH1: a candidate tumor suppressor gene on chromosome 6q24.3 is downregulated in breast cancer. Oncogene. 22:2972–2983.

